# The optoretinogram reveals how human photoreceptors deform in response to light

**DOI:** 10.1101/2020.01.18.911339

**Authors:** Vimal Prabhu Pandiyan, Aiden Maloney Bertelli, James Kuchenbecker, Kevin C. Boyle, Tong Ling, B. Hyle Park, Austin Roorda, Daniel Palanker, Ramkumar Sabesan

## Abstract

Limited accessibility of retinal neurons to electrophysiology on a cellular scale in-vivo has restricted studies of their signaling to in-vitro preparations and animal models. Physiological changes underlying neural activity are mediated by variations in electrical potential that alter the surface tension of the cell membrane. In addition, physiological processes affect concentration of the cell’s constituents that results in variation of osmotic pressure. Both these phenomena affect the neuron’s shape which can be detected using interferometric imaging, thereby enabling non-invasive label-free imaging of physiological activity in-vivo with cellular resolution. Here, we apply high-speed phase-resolved optical coherence tomography in line-field configuration to image the biophysical phenomena associated with phototransduction in human cone photoreceptors in vivo. We demonstrate that individual cones exhibit a biphasic response to light: an early ms-scale fast contraction of the outer segment immediately after the onset of the flash stimulus followed by a gradual (hundreds of ms) expansion. We demonstrate that the contraction can be explained by rapid charge movement accompanying the isomerization of cone opsins, consistent with the early receptor potential observed in the electroretinogram and classical electrophysiology in-vitro. We demonstrate the fidelity of such all-optical recording of light-induced activity in the human retina, namely the optoretinogram, across a range of spatiotemporal scales. This approach incorporates functional evaluation into a routine clinical examination of retinal structure and thus holds enormous potential to serve as a biomarker for early disease diagnosis and monitoring therapeutic efficacy.

## Introduction

Photoreceptors convert light into electrical signals via phototransduction – a process that has been well-characterized using electrophysiological, pharmacological and genetic tools^1^. Since photoreceptors are the primary cells affected in retinal degeneration, and are the target cells in many treatments^2,3^, non-invasive visualization of their physiology at high resolution would be invaluable. In general, the physiological processes associated with neural activity affect the cell’s refractive index^4^ and shape^5–7^, which together alter the light propagation, including changes in light scattering^8,9^, polarization^10^ and optical path length (OPL)^5^. Similarly, cellular deformations accompanying the trans-membrane voltage change during the action potential have been documented in crustacean nerves^11,12^, squid giant axons^13^ and mammalian neurons^7,14,15^. Interferometric imaging of such changes in general, and cellular deformations associated with variations in the cell potential, in particular, offers a non-invasive label-free optical approach to monitor the electrical activity in neurons^6^.

The shape of a cell is determined by the balance of the hydrostatic pressure, membrane tension, and stress exerted by its cytoskeleton^16–18^. Changes in the membrane potential (i.e. de- or hyper-polarization) alter the repulsive forces between the mobile charges within the Debye layer, thereby affecting the membrane tension. Using atomic force microscopy, Zhang et al. demonstrated that HEK293 cells deformed proportionally to the membrane potential change (1 nm per 100 mV)^19^. Using quantitative phase microscopy, Ling et al. demonstrated cellular deformations of up to 3 nm during the action potential in spiking HEK293 cells^6^, and about 1 nm in mammalian neurons^7^. Detecting the ms-fast and nm-scale cellular deformations in transmission is very challenging since the OPL difference between the cells and the surrounding medium scales with the refractive index difference, which is only ≈ 0.02. In reflection, the corresponding OPL changes are about 100 times larger since the refractive index difference does not affect it. OPL changes are further compounded in reflection due to the double-pass interaction of light with tissue. However, weak reflection of light from the tissue boundaries limits the backscattered signal, and hence extracting the nm-scale changes from the shot noise remains quite challenging. This challenge is exaggerated in the living retina, where phase stability is severely affected by eye motion. High-speed and precise image registration is required to retrieve weak signals amidst motion due to respiration, heart beat and fixational eye movements.

Full-field and point-scan phase-resolved optical coherence tomography (pOCT) are powerful vehicles to translate interferometry to the living eye^5,20,21^. Stimulus-induced OPL changes in individual photoreceptors were demonstrated with slower temporal signatures (a few seconds) that align with osmotic changes^22^ related to the intermediary steps and bi-products of phototransduction. Such measurements provided utmost precision in identifying spectral types of cones^21^. A population response averaged across many cones also showed a fast contraction of the outer segments (≈40 nm in ≈5 ms) immediately following stimulus^5,21^. The mechanism behind the fast contractile response in cone photoreceptors remain poorly characterized and understood largely due to the limited temporal resolution available for assessing the events that occur at short ms time scales.

In this article, we introduce a high-speed phase-resolved spectral domain OCT in a line-field configuration, which provides sufficient spatiotemporal resolution to facilitate image registration in living eyes and visualize the rapid biophysical changes during the early stages of phototransduction in individual cones. We demonstrate the stimulus-induced optical response in cones across varying spatial scales, ranging from hundreds of microns to single cells, without and with adaptive optics, respectively. Using high temporal resolution, ranging from milliseconds to microseconds, we demonstrate that the rapid contractile response of the photoreceptors is the optical manifestation of the early receptor potential. We propose a mechanism underlying the electromechanical coupling in photoreceptors based on voltage dependence of the membrane tension.

## Results

Two cyclopleged subjects were imaged at 5 to 7 deg. temporal eccentricity after 3-4 min. dark adaptation using a multi-modal retinal camera, which included a high-speed line-scan spectral domain OCT channel, a line-scan ophthalmoscope (LSO), and a 528±20nm LED source in Maxwellian view for retinal stimulation (photon flux density: 10^6^ – 10^7^ photons/μm^2^)(see Materials and Methods for details). The recorded OCT volumes were reconstructed, registered and segmented for cone outer segment tips (COST) and inner-outer segment junction (ISOS). The temporal evolution of optical phase difference was computed between the cone ISOS and COST (*ϕ*_*COST*_ − *ϕ*_*ISOS*_) to yield a measure of light-induced optical change in the cone outer segments.

### Cone photoreceptors exhibit a repeatable biphasic optical change in response to light stimuli

Spatially patterned light stimuli induce consistent patterned responses, encoded in the phase of the OCT signal detected in the outer retina (Fig 1a-c). Fig 1a shows an LSO cone photoreceptor image of the retina at 7 deg temporal eccentricity without AO, overlaid with a spatially patterned illumination of three horizontal bars drawn to scale. Fig 1b and c show the *en face* OPL between COST and ISOS before (t = −0.29 sec) and after (t = 1.05 sec) the stimulus onset, obtained from the OCT phase difference at the two layers, using the relation *OPL* = (*λ*_*o*_/4*π*) × (*ϕ*_*COST*_ − *ϕ*_*ISOS*_), where λ_o_ = 840 nm.Before the stimulus onset, the phase difference image predominantly denotes the noise floor of the measurement dominated by shot noise, but might also include weak responses due to the imaging light itself activating cone photoreceptors. After stimulus onset, the spatial distribution of OPL replicates the spatially patterned illumination, reaching an amplitude of 200 nm in ≈ 1sec. Note however that the pattern edges are blurred, potentially due to uncorrected aberrations in the stimulus beam introduced through a dilated pupil. Overall, light-activated responses in the cone outer segments have reliable spatial correspondence with the stimulus patterns on the retina.

**Fig. 1.**
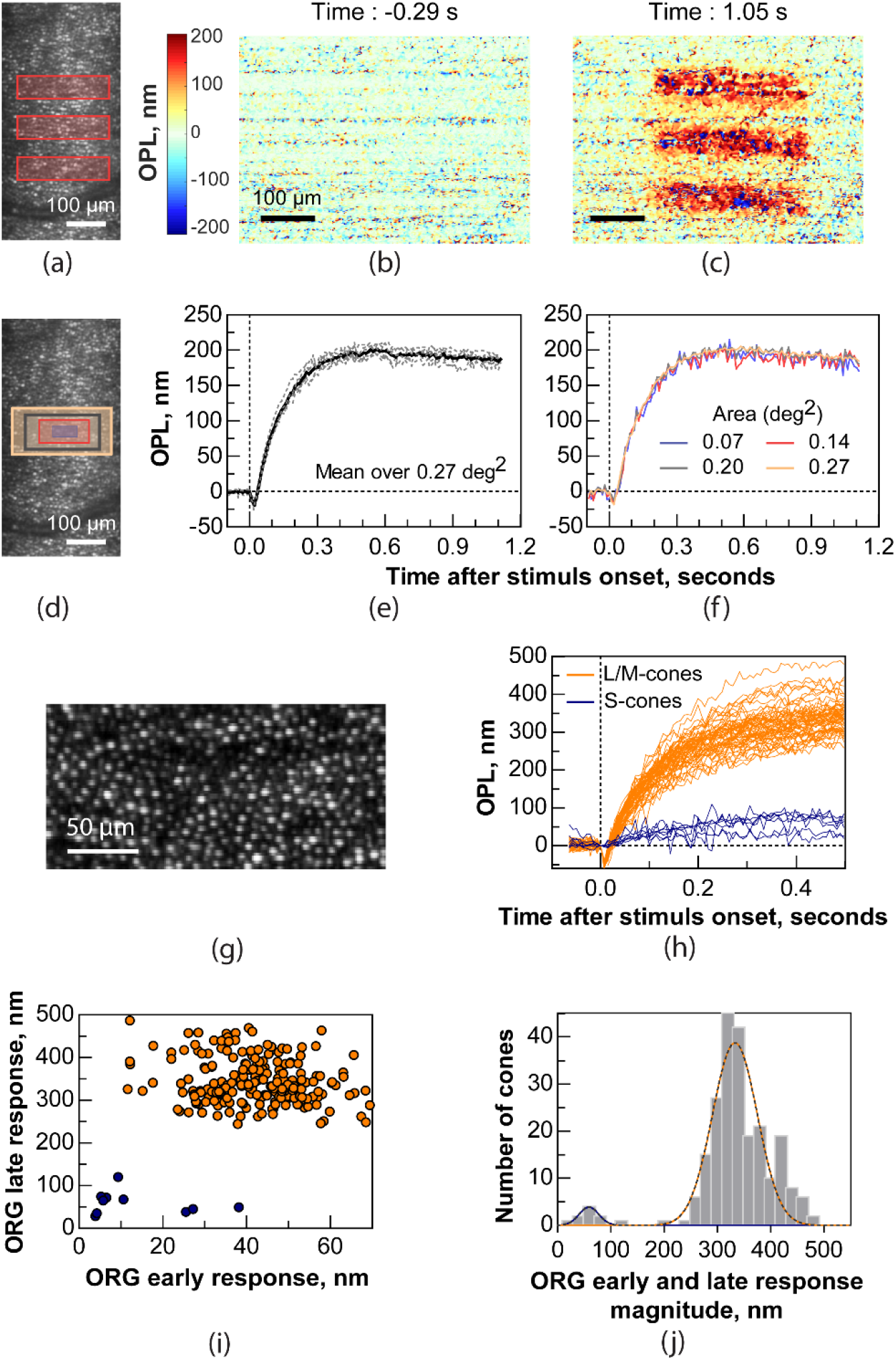
Optoretinography reveals functional activity in cone outer segments across different spatial scales. (a) Illumination pattern drawn to scale over the LSO image. (b,c) The spatial map of OPL change between the ISOS and COST before (b) and after stimulus (c), measured at volume rate of 20 Hz. (d) Rectangles over an LSO retinal image represent the areas over which averages were obtained to plot the ORGs: 0.27 deg^2^ (yellow), 0.20 deg^2^ (gray), 0.14 deg^2^ (red), and 0.07 deg^2^ encompassing ≈10 cones (violet). (e) Repeatability of the response: single ORGs (gray, dotted) and their mean (solid black) for six repeat trials, where phase responses were averaged over 0.27 deg^2^ for 17.9% bleach. (f) Spatial averaging: ORGs over different areas color-coded according to the rectangles in (d). (g) Maximum intensity projection at COST layer with AO reveals individual cone photoreceptors. (h) ORGs for a subset of single cones in (g) demonstrating the response in each cone for 0.3% S-cone bleach and 29.7% average L & M-cone bleach. (i) ORG early and late response amplitudes for each cone in (g). (j) Histogram of the ORG early and late response magnitude, computed as the Euclidean distance from origin of each data point in (i). The histogram is fit with a 2-component Gaussian mixture model (black dotted line) and its component Gaussians are used to distinguish S-cones (blue fit) from L&M-cones (orange fit), on the basis of the different combined response amplitudes attributed to ≈100 times lower bleach in S-cones for the 528nm stimulus compared to L&M-cones. The blue and orange labels in (h) & (i) are obtained from the Gaussian clustering analysis in (j) to denote S- and L&M-cones respectively. The vertical dotted line marks t=0 in (e, f, h) indicate the rising edge of stimulus onset flash. (a-f) are obtained without AO, with 4 mm imaging pupil, at a volume rate of 120 Hz. Fig (g and h) are obtained with AO, for a 6 mm imaging pupil, at a volume rate of 162 Hz.

An LSO cone photoreceptor image of the retina at 7 deg temporal eccentricity taken without AO, overlaid with rectangular stimulus areas ranging from 0.06 to 0.27 deg^2^ is shown in Fig 1d. Six single OPL recordings (gray-dashed lines) after the light onset and their mean (solid line) obtained after averaging phase differences over 0.27 deg^2^ (yellow rectangle in Fig 1d) are shown in Fig 1e. Throughout the article, we will refer to each OPL trace as an optoretinogram (ORG) and more generally, the optical imaging of stimulus-induced activity as optoretinography. As can be seen in Fig. 1e, the magnitude and shape of the ORG was very repeatable across trials; the average variability with and without stimulus was 8 nm and 2.6 nm, respectively. The photoreceptors exhibited an exceptionally reproducible light-driven response: a rapid (<5 ms) reduction in OPL after the stimulus onset, followed by a slower (>1 sec) increase. Based on latency with respect to the stimulus onset, we defined the reduction and increase in OPL as the “early” and “late” ORG responses, respectively.

For uncorrelated noise, averaging N samples in space or time improves the signal-to-noise ratio by 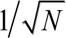. The effect of spatial averaging on the light-activated responses was assessed by determining whether phase differences can be averaged over a small retinal area and still demonstrate reasonable fidelity. Fig 1f shows single OPL traces after the light onset averaged over the areas shown in Fig 1d, demonstrating good consistency in response magnitude and time course for areas as small as that covering 10 cones. The results in Fig 1f represent only single OPL traces, and averaging responses from repeated measurements ought to further improve the signal-to-noise. Results shown in Fig 1a-f were obtained with optimized defocus but without AO.

Next, we focused on the early contraction that has a lower amplitude and faster temporal signature and assessed whether this phenomenon can be reliably resolved on spatial scales as small as individual cones. For this purpose, AO was incorporated to improve lateral resolution and visualize the *en face* cone outer segments (Fig 1g). Fig 1h shows the light-induced OPL for a subset of cones in Fig 1g. Each cone in Fig 1g exhibited the early and late response, with their respective OPL amplitudes differing substantially. Across the 233 and 185 cones tested in Subject 1 and 2, the mean ± std of the early response amplitude was 41.96±10.8 nm and 40.8±14.6 nm, respectively. The early and late response amplitudes for each cone are plotted in Fig 1i. A histogram of the Euclidean distance from the origin denoted the total ORG response amplitude and was subjected to Gaussian mixture model clustering analysis (Fig 1j) to distinguish putative S-cones (4.3% of total cones) from L&M-cones.

### Energy dependence of the early and late cone response

The family of OPL traces (Fig 2) with increasing stimulus (flash) intensity, expressed in photons/μm^2^ and percent bleach, shows that the amplitude of both the late and early responses increases with the stimulus energy (0.1×10^6^ − 6×10^6^ photons/μm^2^ for late response – Fig 2a, 0.04×10^6^ − 2×10^6^ photons/μm^2^ for early response – Fig 2d). The maximum amplitude of the late response vs. flash intensity fits a power function with exponents of 0.5 and 0.6 for subjects 1 and 2, respectively (Fig 2b), growing from 50 to 527 nm. With increasing flash intensity, the magnitude of the early response increased from 5 and 57 nm (Fig 2d). Amplitude of the early response scaled with the stimulus energy according to a logarithmic function for both the subjects, as shown in Fig 2e.

**Fig. 2.**
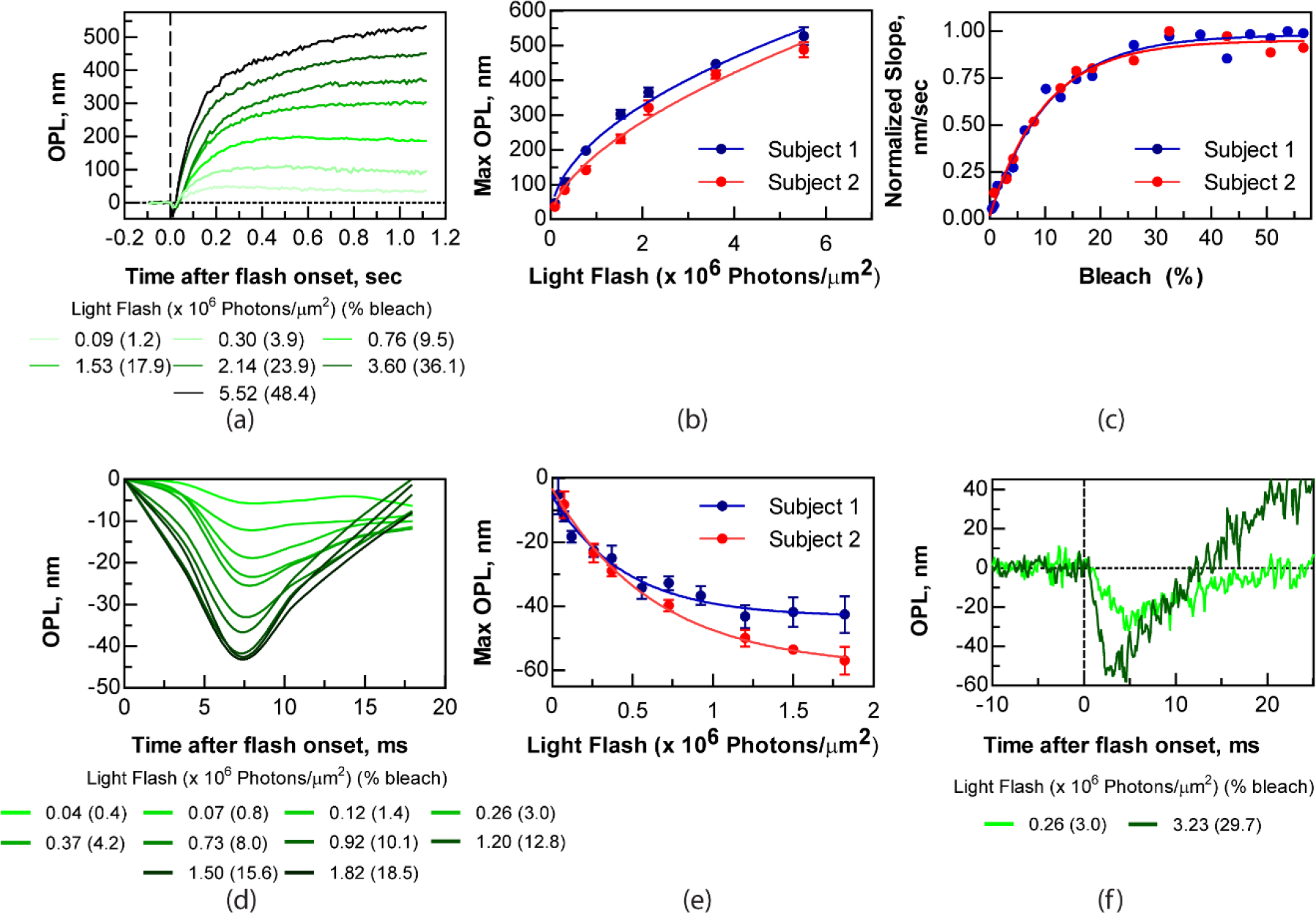
(a) Energy dependence of the optoretinogram. A magnified view of the early response at higher temporal resolution is plotted in d). The maximum OPL change in the late and early responses are shown in (b) and (e), respectively, with their corresponding best fits (solid line in blue and red) for two subjects. Four individual measurements were averaged for all data shown here in fig (a) and (d), error bars denote ± 1 std. dev from the mean. OPL changes for different energy levels are averaged spatially over 0.27 deg^2^. (c) shows the slope of the swelling in the late response for 2 subjects and its best fit. (f) Early response recorded at a high temporal resolution reveals that the onset latency is minimal with respect to the flash onset. Eight individual measurements are averaged in (f). The vertical dotted marks t=0 in (a) and (f) indicates the rising edge of stimulus flash. All the data in these figures was collected without AO and with a 4mm pupil. Volumes rates for (a), (b) is 120 Hz, (c), (d), (e) is 324 Hz and (f) is 32 Hz.

To obtain insight into the rapid initiation of the response and time-to-peak after the stimulus onset, we undertook a modified analysis paradigm that yielded a temporal resolution of 125 μs (see Methods). Fig 2f shows the early response at high temporal resolution for two stimulus intensity levels. The latency of the early response onset and time-to-peak were obtained from bilinear fits to the data (See supplementary material) and estimated to be 0.38 ms (90 % CI : [0.21, 0.51]) & 2.54 ms (CI: [2.43, 2.63]) for 3.0 % bleach and 0.21 ms (90 % CI : [0, 0. 56]) & 4.38 ms (CI: [3.91, 4.76]) for 29.7 % bleach.

We interpret the increase in the optical path length at slow temporal scales (the late response) to be a result of the diffusion of water into the outer segment to maintain osmotic balance during the phototransduction cascade. This was proposed based on observations made in mouse rod outer segments^22^. Based on the observations of Zhang et al., that transducin knockout mice do not show similar osmotic swelling, it follows that the dominant osmolyte implicated in the process must include transducin or its downstream activation stages in the rod phototranduction cascade. There are fundamental differences in the structural architecture of cone and rod outer segments that prevent drawing parallels between them. Chief among them is that rod outer segment disks are disconnected from the plasma membrane, while the membrane in the disks of the cone outer segments is connected to the cilium membranes across the outer segment length. Because in the cone outer segment disks, the membrane surface area in direct contact with the cytosol is larger, they have higher effective osmotic permeability and can achieve osmotic equilibrium faster than rods during phototransduction. For example, at 7.5 % bleach, cone outer segments equilibrate in ≈300 msec, while for 10% bleach, rod osmotic equilibrium takes over 100 seconds^22^. We sought to examine the faster osmotic equilibrium in cones by characterizing the initial slope of the late response. The rate of increase in OPL within 20 ms following the early response peak is plotted as a function of stimulus energy in Fig 2c. The rate of swelling denotes the rate of water entry into the cone outer segments and increases linearly before reaching saturation. The 1/e or 63% of saturation is achieved with ≈15% bleach of the cone opsins. Slope saturation is explained by a complete activation of the phototransduction cascade, leading to an exhaustive consumption and solubilization of the responsible intermediary osmolytes.

A few key features of the fast contractile response provide insight into its mechanistic origin: the latency of the response was short (sub-ms), it peaked earlier with more intense stimuli, the required stimulus intensity was high (sufficient to bleach cone opsins), and the maximum amplitude saturated logarithmically with increasing stimulus intensities. Such short microsecond-scale latencies are typically not associated with the activation stages of phototransduction, even at high stimulus intensities, where typically a few ms elapse between the stimulus onset and initiation of the cascade^23^. Rather, these characteristics of the early response agree well with the early receptor potential (ERP) – a fast electrical signal observed in cone photoreceptors via patch clamp recordings ex-vivo under intense flash stimuli^24–26^. The ERP is attributed to the charge transfer across the cell membrane associated with the conformational change of opsins embedded in the membrane^27^, and is distinct from the later changes in the cell potential due to the closing of ion channels following phototransduction.

Previous observations of the cellular deformations due to changes in the transmembrane potential ^6,19,28^ established a linear relationship between the membrane displacement and cell potential. These are linked by the dependence of the membrane tension on transmembrane voltage (Fig S1b), originating from lateral repulsion of the mobile ions in Debye layer. The fast hyperpolarization of the disk membrane due to charge transfer associated with conformational changes in opsins during isomerization, increases the repulsive forces between charge on the membrane surface, causing the disk membrane to stretch. Since the cell volume is conserved during the ms-short events, expansion of the disks leads to their flattening, as shown schematically in Fig 3a. We established a theoretical model that combines the dynamics of the ERP, the voltage dependent membrane tension, and the mechanics of the folded lipid membrane to describe the contraction of the cone outer segments. Here we outline the key components of the model. A more detailed elaboration of the model appears in a companion article^29^.

**Fig 3.**
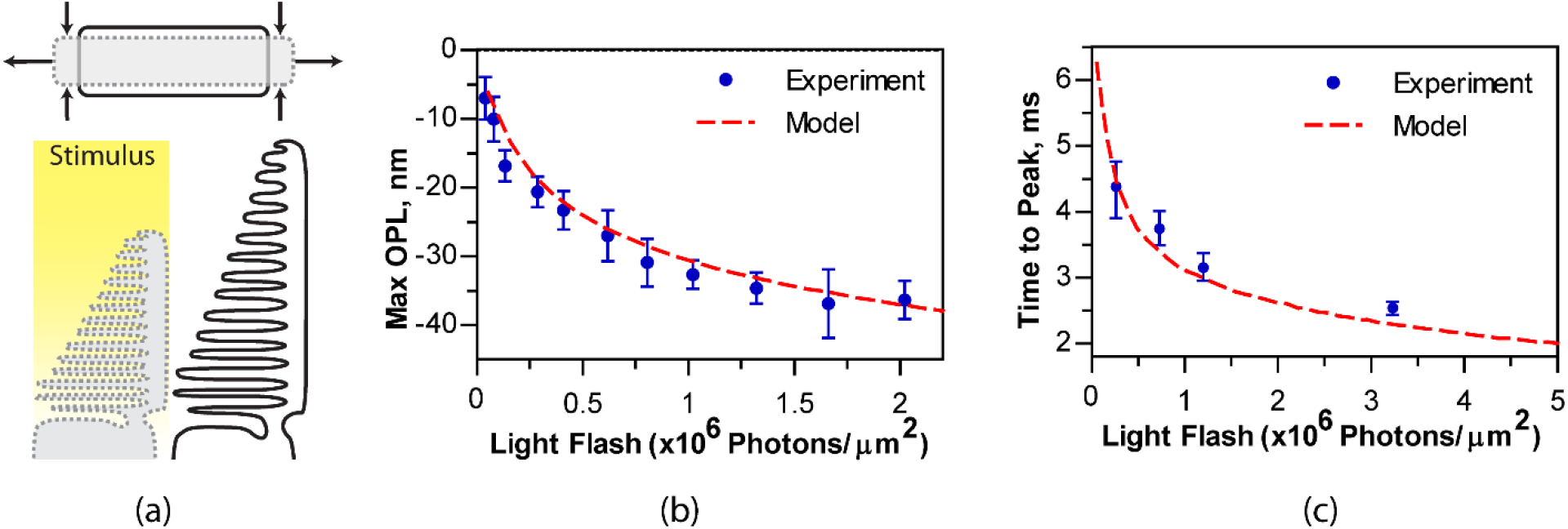
(a) Repulsion of charges in the hyperpolarized disc membrane leads to its expansion, which results in flattening of the disks due to volume conservation. (b) Logarithmic model fit to the maximum amplitude of the early response shown in fig 2(e). (c) Model fit to the time to peak of the early response.

The membrane area expansion coefficient increases with tension as a result of flattening the thermally induced fluctuations^30^. As the membrane stretches, its undulations are reduced and the membrane becomes more resistant to further deformation, as illustrated in Fig. S1b. In a linear approximation of the membrane stiffness scaling with the applied force, the area expansion of the disk *ΔA/A* increases logarithmically with the tension^30^ 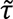:

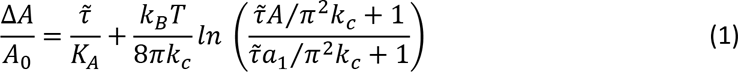

where *A* is the size of the membrane patch (in this case the area of the photoreceptor disk face), *A*_0_ is the area at zero observable tension 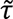, *a*_1_ is the smallest bending feature size (area of a lipid head in a simple lipid bilayer membrane), *k*_*B*_ is the Boltzmann constant, *T* is the temperature, and *k*_*c*_ is the bending modulus. In the case of a membrane densely filled with opsins (typically making up 50% of the membrane area), *a*_1_ is likely not limited by the lipid head width, but rather by the characteristic size of the opsin nanodomains embedded in the disk membrane^31^. With a fixed volume in the disk, the change in thickness of the disk Δ*z*/*z* to accommodate this area expansion is:

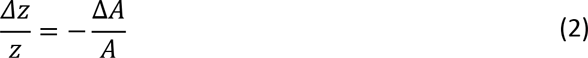

Considering the double pass of light reflected at the bottom of the outer segment, the corresponding change of the OPL in one outer segment disk of height *h* and refractive index *n* is:

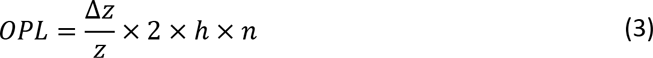

With light intensities well below saturation (<1% bleaching of the opsins), the membrane potential change in the disks does not exceed 2 mV and scales approximately linearly with the number of photons. This corresponds to about 1/100^th^ nm OPL decrease per disk, but in an outer segment of ≈30 μm in height, made up of a stack of hundreds of disks, the overall deformation reaches tens of nm (Fig 3).

The logarithmic relation between the area expansion and membrane tension in equation (1) : *x* ≅ ln(1 + *F*(*x*)) implies an exponential restoring force *F*(*x*) ≅ *e*^*kx*^. By adding a viscous damping term *cẋ*, we arrive at equation of motion with four parameters:

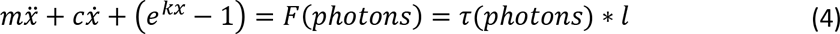

Fitting the parameters (*m*, *c*, *k*, and *l*) to the data yields a very good match to two significant features of the ORG early response. First, the model matches the logarithmic saturation of the amplitude with increasing stimulus intensities, shown in Fig. 3b. Second, it matches the time course of the deformation at different stimulus intensities, specifically the earlier time to peak at higher stimulus intensity, as shown in Fig 3c (also see Fig 2f and Fig S2). The excellent agreement of the model with the experimental results, being consistent in peak deformation and timing across all stimulus intensities with a single set of model parameters (*m*, *c*, *k*, and *l*), indicates that the contraction of the outer segment is an optical signature of the ERP ^29^.

## Discussion

Using interferometric imaging, we quantified the earliest stages of the cone phototransduction in human retina and elaborated the underlying biophysical mechanisms. The optical measurement of such light-induced signals in the retina is termed *optoretinography*, in analogy to classical electroretinography (ERG). ORG terminology was proposed by Mulligan et al^32,33^ in 1994 (although such signals were never detected by them), and adopted first by our group in reference to phase-resolved OCT imaging of the stimulus-induced functional activity in the retina in 2018^34,35^. We restrict our focus here to the photoreceptor component of the ORG.

Recently, phase-resolved OCT in full-field and point-scan configurations were used to measure light-induced responses in cones^5,20,36^. Lack of moving parts and a simple optical layout in full-field approach helps achieving excellent phase stability. However, spatial confocal gating is lost across both lateral dimensions, thus degrading the x-y spatial resolution and contrast. Also, in Fourier domain operation, the frequency-tunable laser source has to be wavelength swept at relatively slow rates to accommodate the 2D sensor acquisition speed, restricting the overall acquisition time per volume (>6ms)^5^. By virtue of high lateral confocality, point-scan operation affords excellent sensitivity, resolution and contrast, at the cost of speed limited by dual-axis scanning^21^. The line-scan protocol employed here provides an optimal trade-off between the speed, resolution and sensitivity for phase-resolved retinal imaging^37^. By losing confocal gating in 1 dimension, it gains improved phase stability along the same dimension and one less moving part (scanner) in the optical system. Because an entire X-Z image is obtained in a single camera snapshot, high phase stability across the entire imaging field of the B-scan is achieved at high temporal resolution. These factors are central to resolving the fast and minute optical changes (ms/nm -scale) associated with mechanical deformations in individual cones.

The time-course of various stages in the phototransduction cascade have been extensively studied^38^. At high stimulus light levels, as used here, that are sufficient to bleach a significant fraction of the cone opsins (≈ 10^5^ – 10^7^ total photoisomerizations per cone), electrophysiological recordings in primate cone photoreceptors are rare. At bright light exposures in rods, the G-protein complex is known to disassociate from the disk membrane and translocate^39^. Zhang et al.^22^ proposed that this causes an osmotic imbalance leading to water uptake into the outer segment. However, G-proteins, specifically, transducin alone could account only for about 1/10^th^ of the entire magnitude of the outer segment swelling in mouse rods, and hence other yet unknown mechanisms or osmolytes might be involved as well. Here, we find that cones differ markedly in their outer segment swelling response to light stimuli, most significantly in the amplitude and rate of saturation. However, the characteristics of the late response demand further study in order to conclusively determine the osmolytes responsible for the effect. Light flashes that bleach several percent of photopigment elicit a similar rate of increase in photocurrent in both rods and cones, consistent with the idea that the rates of activation in early stages of phototransduction – transducin and phosphodiesterase activation – are very similar between rods and cones^23,40^. Therefore, a model of osmotic swelling would need to account for the differences in permeability, outer segment structure and later stages of phototransduction to potentially explain the difference between the swelling magnitude, rate and saturation in rod and cone ORGs.

We observed that the point of inflection, where the early response peaks and the late response initiates, is reached faster at higher stimulus intensities by ≈1.7ms between 3.0% and 29.7% bleach (Figs 2f, 3c). More generally, the rate of change increases with brighter stimuli, i.e. the outer segments undergo a more rapid mechanical deformation, followed by a faster osmotic swelling, compared to that at lower stimulus intensities (Fig 2d and 2f). This is analogous to cone photocurrents measured in vitro, where at higher intensities, both the cone photocurrent amplitude and the initial rate increases, and then eventually saturate when 1-2% of pigment is bleached^23^. For the same bleach level, Hestrin and Korenbrot^23^ also measured a limiting delay for initiation of the cone photocurrent of ≈8 ms after the stimulus onset in salamander retina, substantially longer than the onset of the late response here. Note that much shorter latency (up to 3ms) has been observed in cone-mediated human ERG a-waves^41^, consistent with the latency of the ORG late response observed here: 2.5 ms at 29.7% bleach. Overall, the latency of the initiation of the ORG late response is on a comparable time scale to the onset of cone electrical activity measured in ERGs, though drawing stronger parallels between the temporal features of ORG late response and ERG is reliant on revealing the mechanistic basis of the phototransduction steps implicated in the late response.

The sub-millisecond time scale of the early photoreceptor response is too fast to be affected by the slow osmotic swelling, as the time constant for neuronal swelling is on the order of seconds^42^. It is not likely to be explained by changes of the index of refraction either. Variation in the refractive index due to a conformational change of the opsins upon their isomerization can be estimated using the Kramers-Kronig relation^43^,

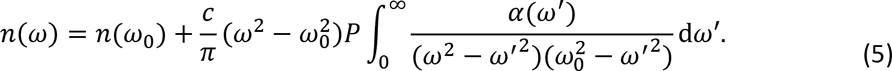

where *n* is the refractive index, *c* is the speed of light in vacuum and *α*(*ω*) is the absorption coefficient at the angular frequency of *ω*. Based on the absorption spectrum shift of rhodopsin upon isomerization^44^, the change in the refractive index at the OCT wavelength (840 nm) would be positive and on the order of 10^-5^, as also measured by Kaplan^45^. This cannot explain the negative change of the OPL in the outer segments by about 10^-3^ during the early response observed here.

The electromechanical deformations of about 40 nm in the outer segments of photoreceptors are much larger than those observed in neurons during an action potential (<2 nm), even though the voltage swing in photoreceptors is much smaller (<10 mV) than during an action potential (≈100 mV). This difference is due to the much softer membranes of the disks and their large number (≈1000) in the outer segments. The disk membranes in photoreceptor outer segments have very low tension since they are formed via a blebbing process and contain no actin cortex^46^. This allows even the slight 2 mV transmembrane potential they experience at 18.5% bleach to yield a ≈0.04 nm deformation on each disk. These minute mechanical changes add up to a very large deformation, proportional mainly to the total number of photoisomerizations in an outer segment containing about 1000 disks.

In conclusion, the optoretinogram enables all-optical label-free monitoring of photoreceptor physiology in humans and has the potential to be applied to other retinal cell types as well. Compared to ERG, the current gold standard for functional evaluation in the clinic, the ORG offers several significant advantages. First, ORGs do not require corneal electrodes. Second, the spatial resolution (testable patch size) is orders of magnitude smaller, and with adaptive optics can be as small as a single cell. Third, ORGs are directly anchored to the OCT structural image, making the spatial localization of the test unambiguous. Finally, while the ORG benefits from adaptive optics, it is not essential, and the ORG can therefore be integrated into conventional OCT imaging systems, enabling not only structural, but also functional evaluation of the retina as part of a routine ophthalmic exam. As such, the ORG has immense potential to serve as an effective biomarker of photoreceptor function that could be used for detecting the earliest manifestations of diseases, their progression, and therapeutic efficacy of treatments.

## Methods

### Experimental setup

A multimodal AO line-scan imager was constructed in free-space to include 3 illumination and 3 detection channels, as shown in Fig 4. A superluminescent diode (λ_o_ = 840 nm, Δλ = 50 nm, 6.2 μm axial resolution in air) source was used as illumination for OCT and for the line-scan ophthalmoscopy (LSO), a 980±10 nm source for wavefront sensing, and a 528±20 nm LED in Maxwellian view for retinal stimulation. A maximum imaging field of 2×2 deg was provided by illuminating a line-field on the retina using a cylindrical lens and scanning it with a 1D galvo-scanner to image volumes. The AO sub-system included a Shack-Hartmann wavefront sensor and deformable mirror (Alpao, DM97-15) incorporated into the optical path. The eye’s pupil, the scanner and the deformable mirror were optically conjugated using lens-based afocal telescopes. An artificial pupil was set to 4 mm and 6 mm for non-AO and AO imaging, respectively, and defocus was optimized using either the deformable mirror or trial lens for non-AO conditions. A lens-based reference arm was used to reduce diffraction due to long distance beam propagation and compensate dispersion at the same time. In detection, a 1200 line-pairs/mm diffraction grating and a high-speed 2D camera formed the spectrometer for OCT. The spatial and spectral resolution (512 × 768 pixels) were optimized in detection by an anamorphic configuration consisting of two cylindrical lenses^47^. The B-scan or M-scan rate was decided by the camera frame rate and varied between 3 – 16.2 kHz in this study. The zeroth-order beam of the grating, which is usually discarded in a traditional spectral-domain OCT system, was used to construct an LSO by placing a focusing lens and a line-scan camera. Because the LSO sensor was optically conjugated to the OCT imaging camera, the enface LSO images were also used to optimize the best focus for imaging. Custom-built software was developed in LabVIEW to synchronize the scanner, frame-grabbers and data acquisition, and allowed real-time feedback to the experimenter for subject alignment via live image visualization. The Nyquist-limited lateral resolution of the system was 2.4 μm, system sensitivity was 92 dB and the phase sensitivity was 4 mrad at an SNR of 50 dB, where 1 mrad ≈ 0.07 nm for λ_o_ = 840 nm. The maximum volume rate used in this study was 324 Hz for 768 × 512 × 50 (*λ* × *x* × *y*) pixels in cases where the temporal resolution of the early response had to be maximized.

**Fig. 4.**
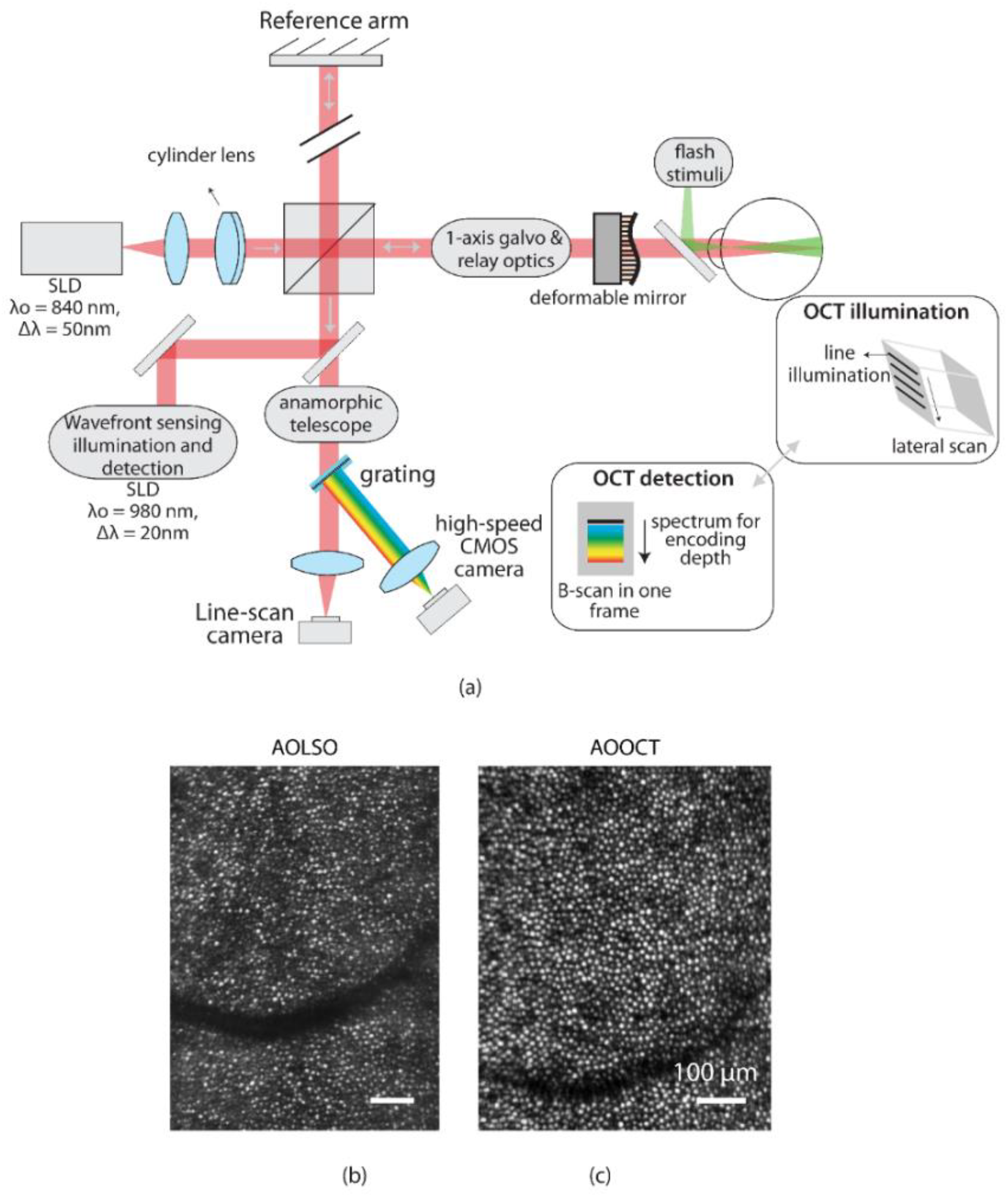
(a) Block diagram of optoretinography system showing key components and features (details in the Methods). AO line-scan ophthalmoscope (b) and OCT images (c) of cone photoreceptors at 7 deg eccentricity. The images are registered, and field flattened to compensate for the Gaussian intensity distribution along the line-dimension.

### Optoretinography protocol

Two cyclopleged subjects free of retinal disease participated in the study. The research was approved by the University of Washington institutional review board and all subjects signed an informed consent before their participation in this study. All procedures involving human subjects were in accordance with the tenets of the Declaration of Helsinki. The subjects were imaged at 5 to 7-deg temporal to the fovea. For ORGs, the subjects were dark adapted for 3-4 min, and OCT volumes were acquired for 1-2 seconds at different volume rates.

After 5 to 20 volumes, a 528±20 nm LED flash illuminated a retinal area of 7 deg^2^ for durations between 0.1 to 70 ms. To optimize temporal resolution and reveal details of the early response, lateral resolution and field of view were sacrificed along the scan dimension, such that a minimum of 2 deg × 0.6 deg (line × scan dimension) were used. To test the performance limits of the system and image the individual cone early responses, a field of view of 0.85 deg and sparse lateral resolution of 4.6 μm were used to achieve maximum volume rates of 162 Hz. Twelve individual measurements were averaged to reveal the early and late responses in individual cones in Fig 1h.

To obtain the latency of the early response, 500 B-scans per volume were recorded at a rate of 16 kHz for a total of 25 volumes in one measurement and eight measurements were averaged for the plots in Fig 2f. The stimulus light onset occurred after the 5^th^ volume. Two B-scans, each taken at 16 kHz, were averaged to obtain a high temporal resolution of 125 μs. Analysis to obtain the latency and peak of the early response is described in main text.

### Image processing

OCT image processing followed conventional techniques. The acquired OCT data were resampled in k-space and Fourier-transformed along the wavenumber dimension to extract complex-valued retinal volumes. For imaging structure, only the absolute value of the complex number was used. Each OCT volume was segmented and en face images were obtained by taking the maximum intensity projections centered at the cone outer segment tips (COST) for visualizing the cone photoreceptors. En face images were registered using a strip-based registration algorithm and averaged ^48^. For optoretinography, the phase analysis followed Hillmann et al.^5^ Each OCT volume was first registered and referenced to the mean of all volumes recorded before the start of stimulus to cancel arbitrary phase noise and set the baseline. From these referenced volumes, a 3-pixel mean of complex values centered at the ISOS and COST was calculated. The phase difference between these two layers was calculated by multiplying the complex conjugate of COST with the ISOS layer and calculating the argument of the resulting product. For strong phase responses that exceed ±*π* radians, phase was unwrapped along the time dimension at each pixel. The change in OPL was calculated by the relation Δ*OPL* = (*λ*_*o*_/4*π*) × (*ϕ*_*COST*_ − *ϕ*_*ISOS*_), where λ_o_ = 840 nm.

## Acknowledgements

Funding for the study was obtained from NIH grants U01EY025501, EY027941, EY029710, EY025501, P30EY001730, Research to Prevent Blindness Career Development Award, Murdock Charitable Trust, Burroughs Welcome Fund Careers at the Scientific Interfaces, Unrestricted grant from the Research to Prevent Blindness. We thank Jay Neitz, Fred Rieke, James Hurley, and Denis Baylor for their input on this work.

VPP, RS, and DP have a commercial interest in a provisional US patent describing the optoretinogram.

## Supplementary material

### 1. Voltage dependent membrane tension

The shape of biological cells is determined by the balance of intracellular hydrostatic pressure, membrane tension, and strain exerted by the cytoskeleton, as illustrated in Figure S1a. The membrane tension has contributions from the lipid bilayer and the lateral repulsion of ions on both the intracellular and extracellular sides of the membrane:

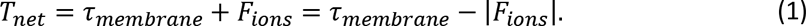

The transmembrane potential changes due to movement of ions across the cell membrane, which affects the charge density in the layers of mobile ions along the inner and outer surfaces of the membrane. This affects the membrane tension due to alteration in lateral repulsion of the ions, and the cell must rebalance the forces via a deformation. The lateral repulsion between the ions can be expressed in terms of the charge density, including the charges accumulated due to membrane voltage V ^1^.

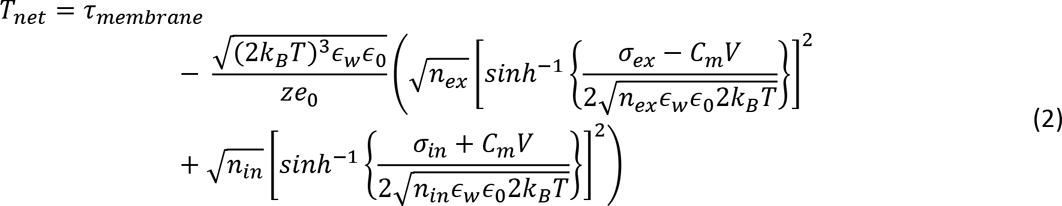

where *σ*_*ex*_ and *σ*_*in*_ are the surface charge densities on the external and internal surface of the cell membrane, *n*_*ex*_ and *n*_*in*_ are the ionic strengths of the extracellular medium and the intracellular cytoplasm, *C*_*m*_ is the specific membrane capacitance (~0.5 μF ∙ cm^−2^), *z* is the valence and *e*_0_ is the elementary charge, *ɛ*_*w*_ is the relative permittivity of water and *ɛ*_0_ is the permittivity of free space, *k*_*B*_ is the Boltzmann constant and *T* is the temperature. The membrane tension increases by about 10 μN ∙ m^-1^ for a 100 mV depolarization^2^ due to decreased repulsive forces between ions in the charge layers. Increased surface tension of the cell membrane leads to deformation of a cell toward minimization of its surface area, i.e. becoming more spherical.

The disk membranes in photoreceptor outer segments have very low tension since they are formed as part of a blebbing process and contain no actin cortex^3^. Due to this lack of structure, they lay flat in stacks that are held together by inter-disk proteins^4^. Any change in *F*_*ions*_ on the photoreceptor disk membrane during illumination affects the membrane tension directly, which can readily stretch in this low-tension state. The low tension allows the membrane to undulate, and even small forces can easily pull the undulations out to allow lateral expansion of the membrane. The membrane area expansion coefficient at low tension is defined by the bending modulus *k*_*c*_ = 0.5×10^−19^ *N* .*m* ^5^, while at high tension, the membrane is mostly flat, and the area expansion modulus *k*_*A*_ = 0.2*N* / *m* dominates^6^. The apparent modulus in-between these two extremes varies as a function of the tension *τ* (Fig S1b):

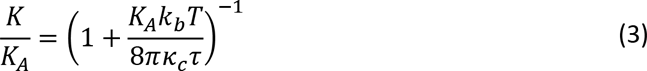

During hyperpolarization of the photoreceptor under illumination, the lateral repulsion of ions in the charge layers *F*_*ions*_ increases and the membrane area will expand. If the cell volume during a few ms of the flash remains constant, widening of the disks will flatten them, leading to shortening of the outer segments.

**Fig. S1.**
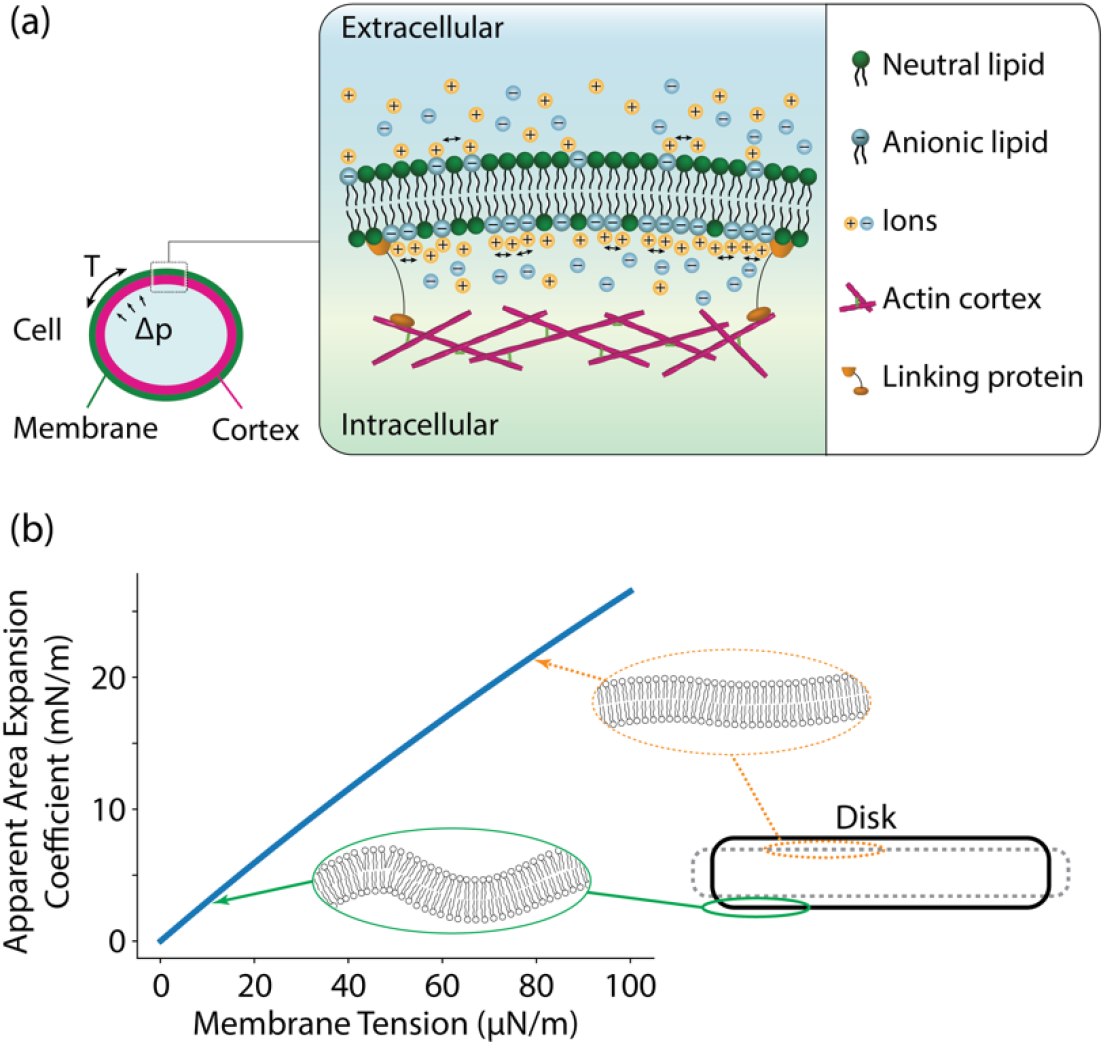
(a) The shape of the cell membrane is defined by the balance of the hydrostatic pressure, membrane tension, and cytoskeleton. Changes in the concentration of mobile ions on either side of the membrane can affect the lateral repulsion of those ions. The resulting change in membrane tension leads to cell deformation. (b) Membrane undulations are flattened under rising tension, thereby increasing its stiffness. Inset shows the reduced membrane undulation at higher tension indicated by arrows.

### 2. Estimating latency and peak of the early response

To determine the latency and the time to peak of the responses from the stimulus onset, we fitted the data with bilinear curves. The point of intersection was obtained as a measure of these values. The fits were bootstrapped in order to obtain the estimates by resampling 90% of the entire dataset across 1000 total iterations. We then computed the 90% confidence interval (5th to 95th percentile; CI) of the estimated latency and the time to peak. For estimating latency, the pre-stimulus slope (baseline activity) was constrained to zero.

**Fig S2:**
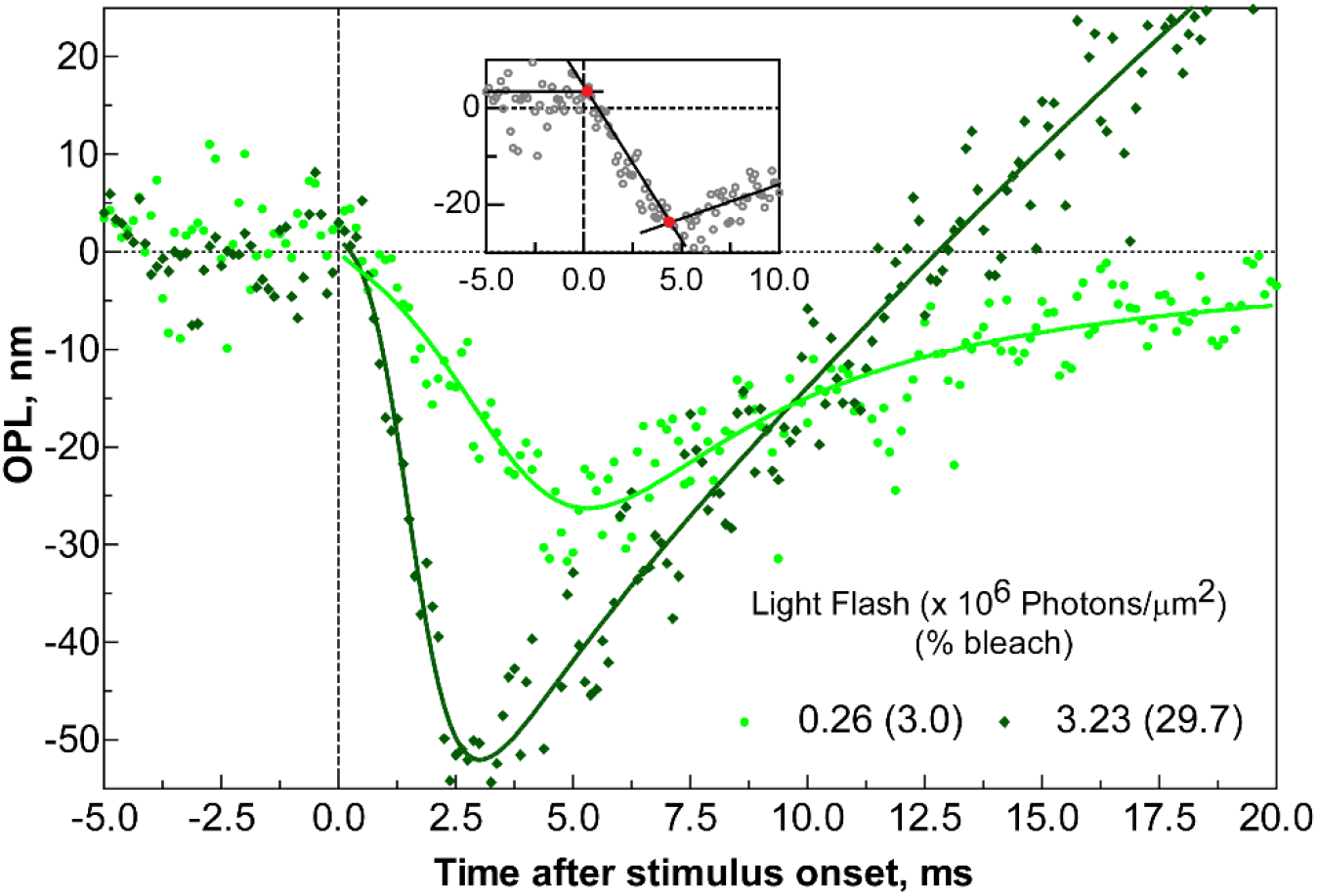
The best fits to the early response at a high temporal resolution (125 μs) for two stimulus intensities (3.0% and 29.7% bleach). Each data point represents a mean of 8 repeated measurements. The inset shows bilinear fits to the data. The first bilinear fit is for the baseline prior to the stimulus and the rising edge of the early response, the intersection of which provides an estimate of the response latency. The second bilinear fit is between the early and late responses, the intersection of which provides an estimate of the response peak. The corresponding values are provided in main text.

## References

1 Arshavsky, V. Y., Lamb, T. D. & Pugh, E. N. Jr., G proteins and phototransduction. Annual review of physiology 64, 153–187, doi:10.1146/annurev.physiol.64.082701.102229 (2002).

2 Lamba, D. A., Gust, J. & Reh, T. A. Transplantation of human embryonic stem cell-derived photoreceptors restores some visual function in Crx-deficient mice. Cell stem cell 4, 73–79, (2009).

3 Bennett, J. et al. Photoreceptor cell rescue in retinal degeneration (rd) mice by in vivo gene therapy. Nature medicine 2, 649 (1996).

4 Foust, A. J. & Rector, D. M. Optically teasing apart neural swelling and depolarization. Neuroscience 145, 887–899, doi:10.1016/j.neuroscience.2006.12.068 (2007).

5 Hillmann, D. et al. In vivo optical imaging of physiological responses to photostimulation in human photoreceptors. Proceedings of the National Academy of Sciences 113, 13138–13143, doi:10.1073/pnas.1606428113 (2016).

6 Ling, T. et al. Full-field interferometric imaging of propagating action potentials. Light: Science & Applications 7, 107, doi:10.1038/s41377-018-0107-9 (2018).

7 Ling, T. et al. How neurons move during action potentials. bioRxiv, 765768, doi:10.1101/765768 (2019).

8 Cohen, L. B., Keynes, R. D. & Hille, B. Light scattering and birefringence changes during nerve activity. Nature 218, 438–441, doi:10.1038/218438a0 (1968).

9 Graf, B. W., Ralston, T. S., Ko, H. J. & Boppart, S. A. Detecting intrinsic scattering changes correlated to neuron action potentials using optical coherence imaging. Opt Express 17, 13447–13457, doi:10.1364/oe.17.013447 (2009).

10 Wang, H. et al. Polarization sensitive optical coherence microscopy for brain imaging. Optics letters 41, 2213–2216, doi:10.1364/OL.41.002213 (2016).

11 Hill, B. C., Schubert, E. D., Nokes, M. A. & Michelson, R. P. Laser interferometer measurement of changes in crayfish axon diameter concurrent with action potential. Science 196, 426–428, doi:10.1126/science.850785 (1977).

12 Iwasa, K. & Tasaki, I. Mechanical changes in squid giant axons associated with production of action potentials. Biochem Biophys Res Commun 95, 1328–1331, doi:10.1016/0006-291x(80)91619-8 (1980).

13 Akkin, T., Landowne, D. & Sivaprakasam, A. Optical coherence tomography phase measurement of transient changes in squid giant axons during activity. The Journal of membrane biology 231, 35–46, doi:10.1007/s00232-009-9202-4 (2009).

14 Kim, G. H., Kosterin, P., Obaid, A. L. & Salzberg, B. M. A mechanical spike accompanies the action potential in Mammalian nerve terminals. Biophysical journal 92, 3122–3129, doi:10.1529/biophysj.106.103754 (2007).

15 Yang, Y. et al. Imaging Action Potential in Single Mammalian Neurons by Tracking the Accompanying Sub-Nanometer Mechanical Motion. ACS nano 12, 4186–4193, doi:10.1021/acsnano.8b00867 (2018).

16 Keren, K. et al. Mechanism of shape determination in motile cells. Nature 453, 475–480, doi:10.1038/nature06952 (2008).

17 Gauthier, N. C., Masters, T. A. & Sheetz, M. P. Mechanical feedback between membrane tension and dynamics. Trends Cell Biol 22, 527–535, doi:10.1016/j.tcb.2012.07.005 (2012).

18 Sens, P. & Plastino, J. Membrane tension and cytoskeleton organization in cell motility. Journal of physics. Condensed matter : an Institute of Physics journal 27, 273103, doi:10.1088/0953-8984/27/27/273103 (2015).

19 Zhang, P.-C., Keleshian, A. M. & Sachs, F. Voltage-induced membrane movement. Nature 413, 428–432 (2001).

20 Azimipour, M., Migacz, J. V., Zawadzki, R. J., Werner, J. S. & Jonnal, R. S. Functional retinal imaging using adaptive optics swept-source OCT at 1.6 MHz. Optica 6, 300–303, doi:10.1364/OPTICA.6.000300 (2019).

21 Zhang, F., Kurokawa, K., Lassoued, A., Crowell, J. A. & Miller, D. T. Cone photoreceptor classification in the living human eye from photostimulation-induced phase dynamics. Proceedings of the National Academy of Sciences of the United States of America 116, 7951–7956, doi:10.1073/pnas.1816360116 (2019).

22 Zhang, P. et al. In vivo optophysiology reveals that G-protein activation triggers osmotic swelling and increased light scattering of rod photoreceptors. Proceedings of the National Academy of Sciences 114, E2937–E2946, doi:10.1073/pnas.1620572114 (2017).

23 Hestrin, S. & Korenbrot, J. I. Activation kinetics of retinal cones and rods: response to intense flashes of light. The Journal of neuroscience : the official journal of the Society for Neuroscience 10, 1967–1973 (1990).

24 Brown, K. T. & Murakami, M. A New Receptor Potential of the Monkey Retina with no Detectable Latency. Nature 201, 626–628, doi:10.1038/201626a0 (1964).

25 Hodgkin, A. L. & Obryan, P. M. Internal recording of the early receptor potential in turtle cones. The Journal of Physiology 267, 737–766, doi:10.1113/jphysiol.1977.sp011836 (1977).

26 Hestrin, S. & Korenbrot, J. Activation kinetics of retinal cones and rods: response to intense flashes of light. The Journal of Neuroscience 10, 1967–1973, doi:10.1523/jneurosci.10-06-01967.1990 (1990).

27 Makino, C. L., Taylor, W. R. & Baylor, D. A. Rapid charge movements and photosensitivity of visual pigments in salamander rods and cones. J Physiol 442, 761–780, doi:10.1113/jphysiol.1991.sp018818 (1991).

28 Oh, S. et al. Label-free imaging of membrane potential using membrane electromotility. Biophysical journal 103, 11–18, doi:10.1016/j.bpj.2012.05.020 (2012).

29 Boyle, K. C. et al. On mechanisms of light-induced deformations in photoreceptors. bioRxiv (2020).

30 Marsh, D. Renormalization of the tension and area expansion modulus in fluid membranes. Biophys J 73, 865–869, doi:10.1016/S0006-3495(97)78119-0 (1997).

31 Rakshit, T., Senapati, S., Sinha, S., Whited, A. M. & Park, P. S. Rhodopsin Forms Nanodomains in Rod Outer Segment Disc Membranes of the Cold-Blooded Xenopus laevis. PloS one 10, e0141114, doi:10.1371/journal.pone.0141114 (2015).

32 Mulligan, J. B., MacLeod, D. I. & Statler, I. C. In search of an optoretinogram. (1994).

33 Mulligan, J. B. The Optoretinogram at 38. (2018).

34 Pandiyan, V. P. et al. Optoretinogram: stimulus-induced optical changes in photoreceptors observed with phase-resolved line-scan OCT. Investigative ophthalmology & visual science 60, 1426–1426 (2019).

35 Sabesan, R., Pandiyan, V. P., Maloney-Bertelli, A., Kuchenbecker, J. A. & Roorda, A. Adaptive optics line-field OCT for high-speed imaging of retinal structure and function. Investigative ophthalmology & visual science 60, 1780–1780 (2019).

36 Zhang, F., Kurokawa, K., Lassoued, A., Crowell, J. A. & Miller, D. T. Cone photoreceptor classification in the living human eye from photostimulation-induced phase dynamics. Proceedings of the National Academy of Sciences, 201816360, doi:10.1073/pnas.1816360116 (2019).

37 Ginner, L. et al. Noniterative digital aberration correction for cellular resolution retinal optical coherence tomography in vivo. Optica 4, 924–931 (2017).

38 Korenbrot, J. I. Speed, sensitivity, and stability of the light response in rod and cone photoreceptors: facts and models. Progress in retinal and eye research 31, 442–466, doi:10.1016/j.preteyeres.2012.05.002 (2012).

39 Sokolov, M. et al. Massive light-driven translocation of transducin between the two major compartments of rod cells: a novel mechanism of light adaptation. Neuron 34, 95–106, doi:10.1016/s0896-6273(02)00636-0 (2002).

40 Cobbs, W. H. & Pugh, E. N. Jr., Kinetics and components of the flash photocurrent of isolated retinal rods of the larval salamander, Ambystoma tigrinum. The Journal of physiology 394, 529–572, doi:10.1113/jphysiol.1987.sp016884 (1987).

41 van Hateren, J. & Lamb, T. The photocurrent response of human cones is fast and monophasic. BMC Neuroscience 7, 34, doi:10.1186/1471-2202-7-34 (2006).

42 Rappaz, B. et al. Measurement of the integral refractive index and dynamic cell morphometry of living cells with digital holographic microscopy. Opt Express 13, 9361–9373, doi:10.1364/opex.13.009361 (2005).

43 Song, W., Zhang, L., Ness, S. & Yi, J. Wavelength-dependent optical properties of melanosomes in retinal pigmented epithelium and their changes with melanin bleaching: a numerical study. Biomedical optics express 8, 3966–3980, doi:10.1364/BOE.8.003966 (2017).

44 Thaker, T. M., Kaya, A. I., Preininger, A. M., Hamm, H. E. & Iverson, T. M. Allosteric mechanisms of G protein-Coupled Receptor signaling: a structural perspective. Methods in molecular biology 796, 133–174, doi:10.1007/978-1-61779-334-9_8 (2012).

45 Kaplan, M. W. Modeling the rod outer segment birefringence change correlated with metarhodopsin II formation. Biophysical journal 38, 237–241, doi:10.1016/S0006-3495(82)84554-2 (1982).

46 Papal, S. et al. The giant spectrin betaV couples the molecular motors to phototransduction and Usher syndrome type I proteins along their trafficking route. Hum Mol Genet 22, 3773–3788, doi:10.1093/hmg/ddt228 (2013).

47 Lu, J., Gu, B., Wang, X. & Zhang, Y. High speed adaptive optics ophthalmoscopy with an anamorphic point spread function. Optics express 26, 14356–14374, doi:10.1364/OE.26.014356 (2018).

48 Stevenson, S. B. & Roorda, A. Correcting for miniature eye movements in high-resolution scanning laser ophthalmoscopy. Vol. 5688 PWB (SPIE, 2005).

## References

1 Zhang, P.-C., Keleshian, A. M. & Sachs, F. Voltage-induced membrane movement. Nature 413, 428–432 (2001).

2 Ling, T. et al. Full-field interferometric imaging of propagating action potentials. Light, science & applications 7, 107, doi:10.1038/s41377-018-0107-9 (2018).

3 Papal, S. et al. The giant spectrin βV couples the molecular motors to phototransduction and Usher syndrome type I proteins along their trafficking route. Human Molecular Genetics 22, 3773–3788, doi:10.1093/hmg/ddt228 (2013).

4 Goldberg, A. F. X., Moritz, O. L. & Williams, D. S. Molecular basis for photoreceptor outer segment architecture. Progress in retinal and eye research 55, 52–81, doi:10.1016/j.preteyeres.2016.05.003 (2016).

5 Hestrin, S. Acylation reactions mediated by purified acetylcholine esterase. Biochimica et biophysica acta 4, 310–321, doi:10.1016/0006-3002(50)90037-0 (1950).

6 Phillips, R. in Physics of Biological Membranes (eds Patricia Bassereau & Pierre Sens) 73–105 (Springer International Publishing, 2018).

